# Model for chromosome congression based on motor proteins in microtubule overlaps

**DOI:** 10.1101/2024.05.10.593475

**Authors:** Ivan Sigmund, Domagoj Božan, Ivana Šarić, Nenad Pavin

## Abstract

Mitotic spindle, a micromachine composed of microtubules and associated proteins, plays a pivotal role in ensuring the accurate segregation of chromosomes. During spindle assembly, initially randomly distributed chromosomes are transported toward the equatorial plate and experiments suggest that several competing mechanisms can contribute to this process of chromosome congression. However, a systematic theoretical study of forces relevant to chromosome congression is still lacking. Here we show, by introducing a physical model, that length-dependent forces generated by motor proteins transport chromosomes toward the spindle equator. Passive crosslinkers, on the other hand, can generate off-centering forces that impair chromosome congression. Our mean-field approach also reveals that stable points can exist in the vicinity of spindle poles, in addition to the one in the center, and thus provides an explanation for erroneous spindles with polar chromosomes. Taken together, our study provides a comprehensive approach to understanding how different spindle components interact with each other and generate forces that drive chromosome congression.

## I. INTRODUCTION

The ability of the cell to replicate itself by cell division is one of the basic principles of life. To ensure proper segregation of duplicated genetic material between two daughter cells, the cell arranges chromosomes in space and time. At the beginning of mitosis, the cell exerts forces that transport the chromosomes from initial random positions toward the future division plane in a process termed congression [1–5], as depicted in Fig. 1. Once congression is completed and chromosomes are aligned to the metaphase plane, mitosis proceeds by initiating the segregation of chromosomes. Thus, a precise coordination of forces in space and time is required for the arrangement of chromosomes during cell division.

**FIG 1.**
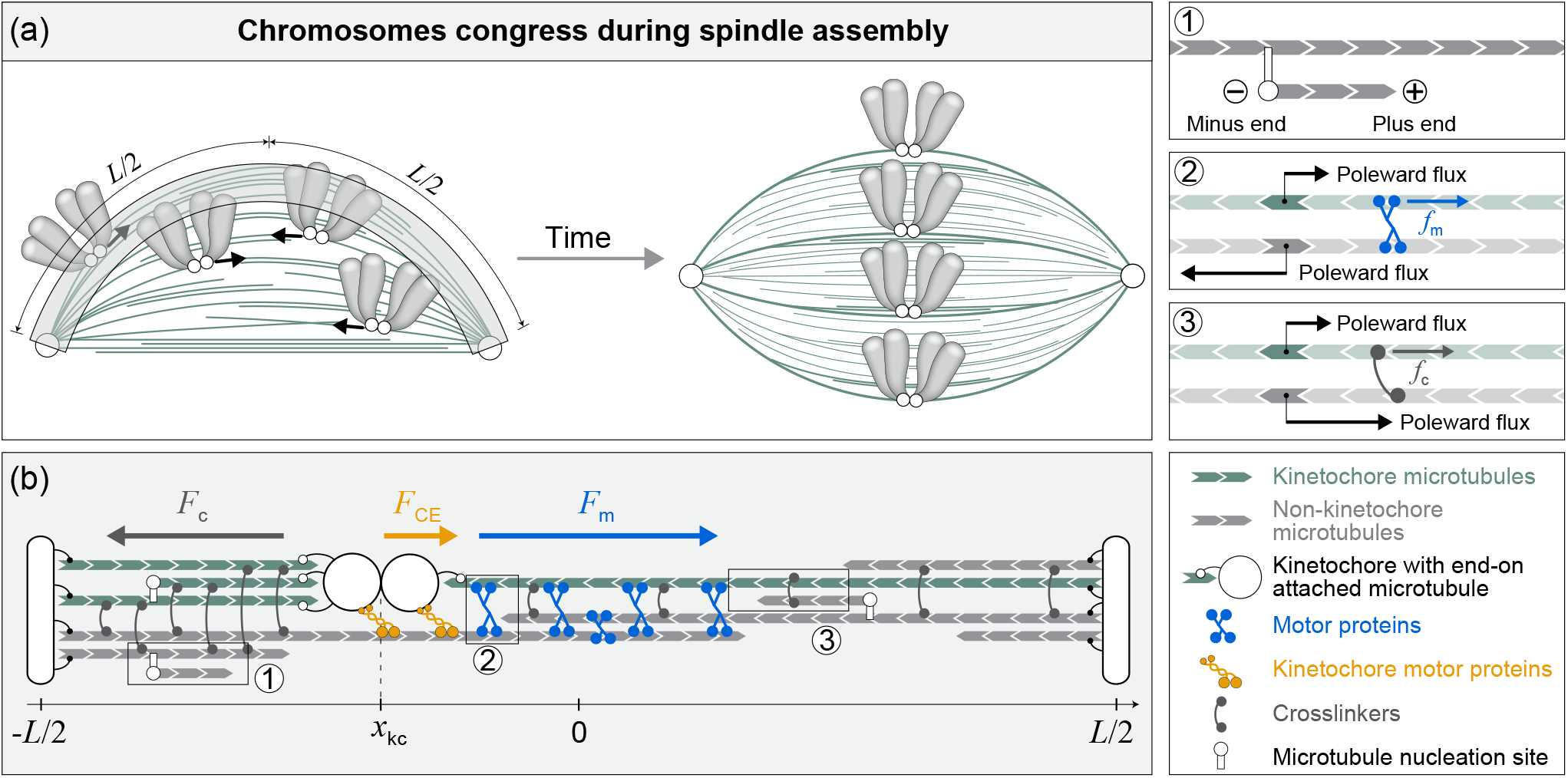
Model for chromosome congression. (a) Schematic representation of chromosome congression during spindle assembly, where the chromosomes move from initial random positions (left scheme) to the spindle equator (right scheme). MTs (green lines) extend between spindle poles (white circles) and interact with kinetochores (white circles at the chromosomes). Arrows represent chromosome movement direction. (b) Geometry of the one-dimensional model with spindle center at location *x* = 0, two poles at locations −*L/*2 and *L/*2, and kinetochores at *x*_kc_. MTs are attached to kinetochores or not. They extend from spindle poles or nucleation sites at pre-existing MTs. Motor proteins link antiparallel MTs and exert force *F*_m_, while passive crosslinkers link parallel MTs and exert force *F*_c_ on the kinetochores. Kinetochore motor proteins link kinetochores to MTs, exerting force *F*_CE_. Individual parts of the scheme are explained in the legend on the right. (inset 1) Enlarged segment showing MT nucleated along a pre-existing MT. The more dynamic plus end is denoted with plus sign and the less dynamic minus end is denoted with a minus sign. (inset 2) Motor protein links kinetochore MT with its antiparallel neighbor and exerts force *f*_m_ on the kinetochore MT. The arrows denote direction of poleward flux of respective MTs. (inset 3) Passive crosslinker links kinetochore MT with its parallel neighbor and exerts the force *f*_c_ on the kinetochore MT. The arrows denote direction of poleward flux of respective MTs.

The mitotic spindle is a bipolar micromachine composed of microtubules (MTs) and associated proteins that self-assembles at the beginning of mitosis and generates the forces responsible for the chromosome positioning [5–8]. Spindle assembly relies on dynamic properties of MTs, including polymerization, depolymerization and stochastic transitions between these two states [9]. Spindle MTs can extend from spindle poles and interact with chromosomes via kinetochores, termed kinetochore MTs [10]. Spindle MTs that extend from opposite poles and interdigitate, enabling motor and non-motor proteins to crosslink them, and thus perform various biological functions [11–17], are termed non-kinetochore MTs. These interactions between MTs and chromosomes, mediated by motor proteins, drive chromosome congression.

Forces directed toward the spindle midplane are crucial for chromosome assembly, and various mechanisms have been proposed to explain how these forces arise [18]. Experiments show that MTs extending from poles exert a force that pushes chromosomes away from the pole, known as polar ejection force, and that this force is generated by motor proteins at chromosome arms [19–23]. Forces oriented toward the spindle midplane are also exerted by plus-end-directed motor proteins at kinetochores, such as CENP-E/kinesin-7, which move along non-kinetochore MTs [24, 25]. A recent study has shown that chromosome congression relies on end-on kinetochore-MT attachments and in their absence, chromosomes do not congress [26]. Pulling forces on kinetochores are generated by depolymerization of end-on attached MTs, as shown *in vitro* and *in vivo* [27–30]. Because forces at sister kinetochores are oriented toward opposite spindle poles, the force toward the center in this tug-of-war situation is generated when longer MTs exert greater forces than the shorter ones. The MT length-dependent forces is produced by accumulation of greater number of kinesin-8 motor proteins at the plus end, where they promote MT depolymerization and consequently increase pulling forces at the kinetochore to which they are attached [31–34]. Alternatively, the longer MT can also generate greater force through motor proteins that accumulate along it in a length-dependent manner and generate faster poleward flux [35]. All these studies suggest that different mechanisms contribute to chromosome congression at the same time.

Theoretical models that describe spindle mechanics give quantitative insights into key processes of mitosis. An important process relevant to mitosis is the formation and stability maintenance of antiparallel MT bundles, which has been studied for a pair of antiparallel MTs [36, 37], as well as for MTs emanating from two poles or along pre-existing MTs [38, 39]. Models for spindle formation also rely on MT reorientation that leads to MT alignment, which is driven by motor proteins and passive crosslinkers [40, 41]. Several theoretical models have studied how MTs and kinetochores explore the space to get into the proximity of each other and subsequently form kinetochore MT attachments, including biorientation [42–48]. In order to study chromosome positioning, in yeasts, the models explored contribution of length and tension dependent regulation of MT dynamics [49– 53], whereas in higher eukaryotes, the models incorporate polar ejection forces and forces generated by kinetochore motor proteins [54–58]. Recently, we have shown that length-dependent MT poleward flux generates centering forces at the kinetochores during metaphase [35]. However, important mechanisms that could drive chromosome congression, such as MT length-dependent force and its cooperation with kinetochore motor proteins have not been explored so far.

In this paper, we introduce a one-dimensional model describing forces that drive chromosome congression in human spindles. These forces are generated by motor proteins and passive crosslinkers that are attached to kinetochore MTs and drive poleward flux of these MTs. To determine the numbers of motor proteins and crosslinkers, we calculate MT distributions from their dynamic properties. The key process behind the centering mechanism is the competition between length-dependent pulling forces and the number of MTs that attach to kinetochores. By using parameters that are relevant for human cells, we find that motor proteins lead to congression times comparable to experimentally measured values. Passive crosslinkers, on the other hand, can generate off-centering forces that impair chromosome congression, but in competition with motor forces they are weaker and do not affect centering. Kinetochore-associated motor proteins generate a centering force that works together with these forces. Thus, our theory presents a methodical framework for understanding the interactions among diverse spindle components and the forces they generate to transport chromosomes during spindle assembly.

## II. THE MODEL

In order to study chromosome congression, we introduce a one-dimensional model that describes the distribution of MTs and the forces that arise in the interaction of MTs with kinetochores (Fig 1). These forces are caused by the flux of MTs, which depends on the length of the MTs, as a consequence of the accumulation of motor proteins, such as Eg5/kinesin-5 [59–61], and crosslinker proteins, such as NuMA [62], on the MTs. The model also includes forces generated by plus-end-directed kinetochore motor proteins, such as CENP-E. The key part of our model is MT length-dependent force, where longer overlaps of antiparallel MTs accumulate a greater number of motor proteins. For this reason, longer MTs that are attached to kinetochore from the farther pole side will generate a greater force than those from the near pole side, thus centering the chromosome.

In order to evaluate this force, we need to calculate the distributions of MTs. Because MTs in human spindles are predominantly aligned with each other [8], we use a one-dimensional approach in which we consider MT distributions with respect to the line parallel to the MTs. We separately consider the MTs that are attached to kinetochores and those that are not, where each of them is extending from the pole or pre-existing MTs. In addition, MTs that are not attached to kinetochores are growing or pausing. Because MT distributions are dynamically generated, we calculate them by taking into account known MT properties, including nucleation, growth, pausing, and catastrophe.

### A. Forces relevant for chromosome congression

In our one-dimensional approach, the position along the spindle is denoted *x* and can take values between −*L/*2 and *L/*2, representing the left and right poles, respectively (Fig. 1). Furthermore, *x* = 0 denotes the position of the spindle center. To calculate how the position of the center of mass of the sister kinetochores, *x*_kc_, changes in time, *t*, we consider the force balance at the kinetochores:

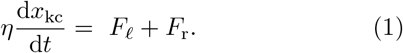

Here, *η* denotes the cytoplasmic drag friction coefficient of the chromosome, and *F*_*ℓ*,r_ denotes the forces at the kinetochore. The indices *ℓ* and r denote the left and right sister kinetochores, respectively.

Forces on the kinetochores arise from the MT plus-end interaction with the kinetochores or from the interaction with kinetochore motor proteins that link kinetochores to MTs laterally,

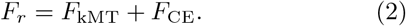

Here, the interaction between the right kinetochore and *N*_k_ MTs attached to it is calculated as and crosslinkers on the *i*-th kinetochore MT are denoted 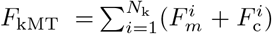. The forces exerted by motor proteins and crosslinkers on the i-th kinetochore MT are denoted 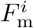 and 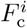, respectively. The force exerted by kinetochore motor proteins is denoted *F*_CE_. Note that here and in the rest of this section we consider forces exerted on the right-hand side only, whereas the complete description of the model is provided in Appendix A.

The forces exerted by motor proteins and crosslinkers attached to the *i*-th kinetochore MT are given as

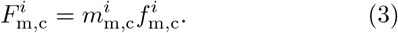

Here, the number of motor proteins and crosslinkers is denoted as 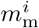, and 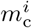, respectively. The average forces exerted by these motor proteins and crosslinkers are denoted 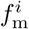 and 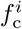, respectively. We calculate the number of motor proteins and crosslinkers attached to the *i*-th kinetochore MT of length *l*^*i*^ as

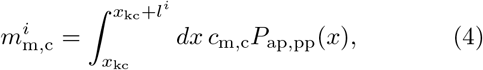

where the integration limits, and consequently number of motor proteins and crosslinkers, depend on the kinetochore position. Here, we take into account that the distribution of motor proteins and crosslinkers depend on the probability of binding to antiparallel neighboring MTs *P*_ap_(*x*) and parallel neighboring MT *P*_pp_(*x*), respectively. Linear densities of motor proteins and crosslinkers are denoted *c*_m_ and *c*_c_, respectively. The fraction of motor proteins that bind to the kinetochore MT and a neighboring antiparallel MT is proportional to the probability of finding an antiparallel neighbor at that position, 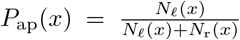. Similarly, the fraction of passive crosslinkers that bind to the kinetochore MT and a neighboring parallel MT is proportional to the probability of finding a parallel neighbor at that position, 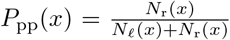. Here *N*_*ℓ*_ and *N*_r_ denote the numbers of left and right non-kinetochore MTs, respectively. We also define the lengths of parallel and antiparallel overlaps with kinetochore MT of length *l*^*i*^ as 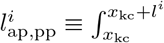. As linear densities of motor proteins and crosslinkers are constant, Eq. (4) simplifies to 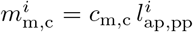. By using these overlap lengths we calculate the average overlaps for the kinetochore MT bundle composed of *N*_k_ MTs as

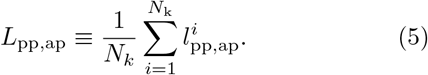

We calculate the velocity of motor proteins from the force-velocity relationship *v*_m_ = *v*_0_(1 − *f*_m_*/f*_0_). Here, *f*_m_ is a load force that opposes motor movement, *f*_0_ is the stall force of the motor, and *v*_0_ is the velocity of the motor protein without a load. The motor velocity is equal to the relative sliding velocity of antiparallel MTs, where 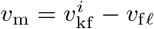 describes the case in which kinetochore MT and the associated non-kinetochore MT have poleward flux 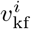 and *v*_f*ℓ*_, respectively. On the other hand, crosslinkers cannot generate an active force between MTs, but if MTs slide between each other, the crosslinkers oppose this movement by exerting a damping force. In our model, the damping force of a crosslinker that connects parallel MTs is given as *f*_*c*_ = −*ξ*_c_*v*_c_, where *ξ*_c_ denotes the viscous friction coefficient. The relative sliding velocity of the kinetochore MT and the associated non-kinetochore parallel MT is given as 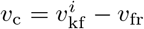.

In order to quantify the forces on the kinetochore MTs, we impose that all kinetochore MTs of the same bundle slide with the same velocity, 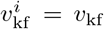, and consequently all the forces in the same bundle have the same value, 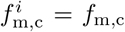. This constraint, which dramatically reduces the number of unknowns, is plausible because MTs within the same kinetochore MT bundle are connected by passive crosslinking proteins. In this limit, the force between the MTs and the kinetochore simplifies to

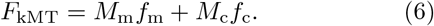

The number of motor proteins and crosslinkers on the kinetochore MT bundle is given as 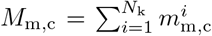 which in combination with Eqs. (4) and (5) yields

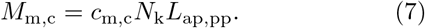

This equation is the core of the length-dependent force at the kinetochore. Note that the average overlap lengths depend on the position of the kinetochore, which in turn affects its positioning. We also use a constant flux velocity for non-kinetochore MTs, *v*_f*ℓ*,fr_ = ∓*v*_0_*/*2, based on results from [35].

The interaction between a kinetochore and the MTs attached to it is given by a relationship between the pulling force exerted by the kinetochore and the growth velocity of the MTs [63]. We simplify this relationship with a linear expression 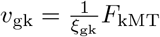. Here, the growth velocity of the kinetochore MTs, *v*_gk_, is calculated from the relative movement of kinetochore MTs, which flux with the poleward velocity *v*_kf_, with respect to the kinetochore, which moves with velocity *v*_kc_ ≡ *dx*_kc_*/dt*, resulting in *v*_gk_ = *v*_kf_ − *v*_kc_. The effective friction coefficient is denoted *ξ*_gk_.

The model also includes kinetochore motor proteins which exert forces by moving along non-kinetochore MTs in a plus-end-directed manner. We describe the force exerted by *N*_CE_ kinetochore motor proteins as

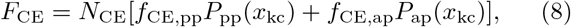

where the direction and magnitude of the forces exerted by kinetochore motor proteins depend on MT orientation. For the right sister kinetochore, described by Eq. (8), the forces exerted toward the left and right by kinetochore motor proteins are denoted *f*_CE,pp_ and *f*_CE,ap_, respectively. These forces are calculated from the force velocity relationships 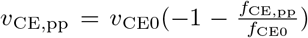 and 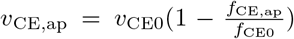. The velocity of kinetochore motor proteins is calculated as the relative velocity of the kinetochore with respect to the associated non-kinetochore MT, *v*_CE,pp_ = (*v*_kc_ − *v*_fr_) and *v*_CE,ap_ = (*v*_kc_ −*v*_f*ℓ*_). Here, *f*_CE0_ denotes the stall force of the kinetochore motor proteins and *v*_CE0_ the velocity without load.

### B. Distributions of kinetochore and non-kinetochore microtubules

In order to calculate the number of MTs, overlap lengths as well as the probabilities *P*_ap,pp_ used in Eqs. (7) and (8), we need to evaluate the MT distributions, which we obtain using a mean-field approach. This approach neglects fluctuations and thus it is suitable for systems with large number of MTs. In our case, it is adequate for description of non-kinetochore MTs, whose number in metaphase is around 5000, whereas it provides an estimate for kinetochore MTs, whose number is around 10, as measured in Ref. [8]. For these reasons, the mean-field approach is relevant for describing typical kinetochore movement, whereas stochastic simulations are more appropriate for studying the variability in the movement of individual kinetochores, as in Refs. [47, 48, 58].

Our model describes growing and pausing MTs, nucleated either on the pole or along pre-existing MTs. The densities of these four distributions are denoted *n*_p_(*l*), *ñ*_p_(*l*), *ρ*_n_(*l, x*), and 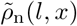, respectively. These densities determine the number of MTs of length *l* extending from the pole, *dN*_p_ = (*n*_p_ + *ñ*_p_)*dl*, and the number of MTs of length *l* extending from a nucleation site at position *x*, 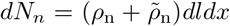. Following approaches from previous studies [64–66], the MT densities are calculated from their dynamic properties using transport equations. For MTs that extend from the pole, the equations are:

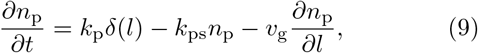

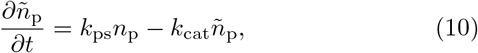

and for those that extend from pre-existing MTs, the equations are:

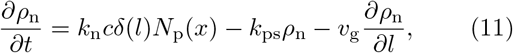

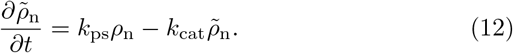

In Eq. (9) the first term on the right-hand side describes MT nucleation at the pole that occurs with rate *k*_p_, where the Dirac delta function, *δ*(*l*), ensures that nucleated MTs have zero length. The second term describes the MT switch from growing to pausing, which occurs at rate *k*_ps_. The last term describes MT growth, in which the drift velocity is an effective MT growth velocity *v*_g_. The pausing MTs extending from the pole are described by Eq. (10), and they disappear upon catastrophe with rate *k*_cat_. The number of pre-existing MTs extending from the pole includes pausing and growing MTs and it is calculated as 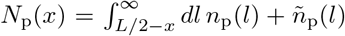. In Eq. (11) the first term describes MT nucleation along pre-existing MTs extending from the pole with nucleation rate multiplied by the density of nucleation sites along a single MT, *k*_n_*c*. The pausing MTs nucleated along pre-existing MTs is described by Eq. (12).

Growing MTs can attach to kinetochores when the MT plus-end is close to the kinetochore. The number of kinetochore MTs extending from the pole is calculated as

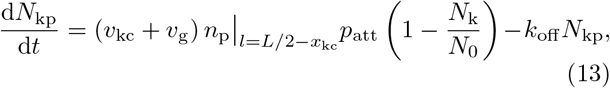

whereas the density of kinetochore MTs nucleated along pre-existing MTs is a function of MT length and is calculated as

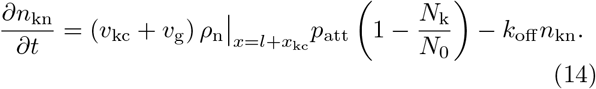

In these two equations, the first term on the right-hand side is the rate of MT attachment to kinetochores, which is proportional to the current of MT plus-ends at the kinetochore position. The current is calculated as the velocity of the MT plus ends with respect to the kinetochore multiplied by the density of plus-ends of non-kinetochore MTs. The attachment probability to an unoccupied kinetochore site is denoted *p*_att_. A fraction of unoccupied attachment sites is calculated as the number of unoccupied attachment sites, *N*_0_ − *N*_k_, divided by the number of attachment sites, *N*_0_. MTs detach from kinetochores at a rate *k*_off_ .

The number of kinetochore MTs is given as:

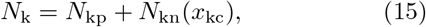

where the number of kinetochore MTs nucleated along pre-existing MTs and reaching position *x* is calculated as

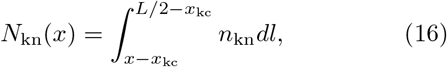

which is valid for *x* between *x*_kc_ and *L/*2. The overlap lengths, *L*_ap_ and *L*_pp_, are calculated as

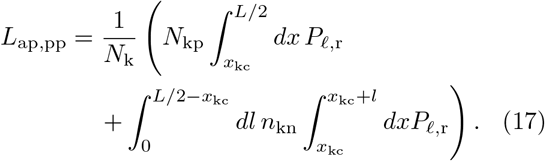

In this mean field approach the sum over individual kinetochore MTs from Eq. (5) is split into two parts. The first part describes kinetochore MTs that extend from the pole to the kinetochore and all have the same length. The second part of the sum describes kinetochore MTs nucleated along pre-existing MTs.

In this section, we have presented a 1D model for chromosome congression, which includes MT dynamics and forces exerted by motor proteins, crosslinkers and kinetochore motor proteins. We aim to calculate the kinetochore velocity, *v*_kc_, as it is a key indicator of the vitality of the congression and the stability of the metaphase plate. Using the equations for the balance of forces on the kinetochores, Eq. (1) and Eq. (2), together with the expressions for kinetochore MT growth velocities, Eq. (A5), average motor protein forces, Eq. (A6), average crosslinker forces, Eq. (A7), and average kinetochore motor protein forces, Eq. (A8), we can calculate the kinetochore velocity, *v*_kc_, as provided in Appendix B. The resulting formula for the kinetochore velocity as a function of the numbers of motor proteins, crosslinkers and kinetochore motor proteins is:

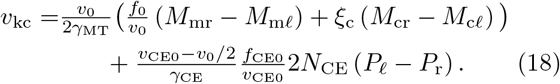

Here, 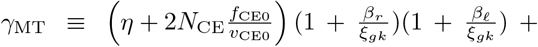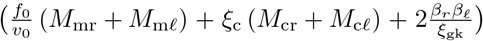 is a short-hand notation for a positive function of *x*_kc_, and it represents an effective drag coefficient for the interaction between MTs and the kinetochore. The shorthand notation 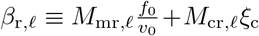 is the sum of motor proteins and crosslinkers on the right or left side, respectively. The shorthand notation for the effective drag coefficient from the interactions between the kinetochores and the kinetochore motor proteins is 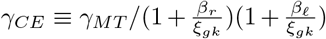 Furthermore, *P*_pp_ and *P*_ap_ have been replaced with 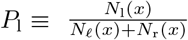 and 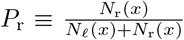, since the same equations describe kinetochore motor proteins on both kinetochores. The terms in parentheses of Eq. (18) show that the direction of the velocity is dictated by the differences in the total number of motor proteins, crosslinkers, and kinetochore motor proteins on the right and left side, each contributing to the velocity with different magnitudes.

## III. RESULTS

In our model, the movements of the kinetochores are governed by a combination of forces exerted by motor proteins and passive crosslinkers on the kinetochore MTs, as well as kinetochore motor proteins (see Eq. (18)). Additionally, distributions of these proteins depend on the geometry of the system, including the number of MTs and their average length, and thus getting a deeper insight into the contributions of different proteins is challenging. Therefore, we study the contributions of each protein separately, as well as concurrently. In this way, we gain a comprehensive understanding of how they work together to ensure proper chromosome congression.

We solve the MT distribution equations in a steadystate limit (see Appendix C), which is reasonable due to the difference in time scales. For non-kinetochore MTs, the MT growth time, 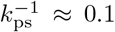 min, is much shorter than the time needed for congression, usually 5-10 min [26, 75]. We also calculate the kinetochore MT distributions in a steady-state limit (see Appendix D), because the kinetochore MT detachment time, 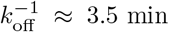 [70], is shorter than the congression time. We selected the model parameters based on recent *in vitro* and *in vivo* measurements. Table I shows the MT and kinetochore parameters, and Table II shows the motor protein, crosslinker and kinetochore motor protein related parameters.

**TABLE I.** Microtubule and kinetochore parameters.

| Parameter | Value | Description |
| --- | --- | --- |
| $L$ | 12 $\mu\text{m}$ | Pole to pole distance [67] |
| $l_0$ | 2.6 $\mu\text{m}$ | MT length<sup>a</sup> |
| $k_{\text{ps}}$ | 11 $\text{min}^{-1}$ | MT pause rate<sup>b</sup> |
| $v_g$ | $k_{\text{ps}}l_0 \approx 28 \mu\text{m}/\text{min}$ | MT growth velocity |
| $k_{\text{cat}}$ | 0.6 $\text{min}^{-1}$ | MT catastrophe rate<sup>c</sup> |
| $k_p$ | 600 $\text{min}^{-1}$ | MT nucleation at poles<sup>d,h</sup> |
| $k_{nc}$ | 0.5 ($\mu\text{m min}$)<sup>-1</sup> | MT nucleation along MTs<sup>e</sup> |
| $N_0$ | 15 | Max. number of kMTs [69] |
| $p_{\text{att}}$ | 0.02 | kMT attachment probability<sup>f,h</sup> |
| $k_{\text{off}}$ | 0.35 $\text{min}^{-1}$ | kMT detachment rate [70] |
| $\xi_{\text{gk}}$ | 300 pN min / $\mu\text{m}$ | Effective friction coefficient<sup>g</sup> |
| $\eta$ | 0.08 pN min / $\mu\text{m}$ | Cytoplasmic drag [71] |

<sup>a</sup> Estimated using electron microscopy data [8] which measured a spindle length of 9.48 $\mu\text{m}$ and average non-kinetochore MT length of 2.07 $\mu\text{m}$ . By scaling to spindle length of 12 $\mu\text{m}$ , we calculate average MT length of 2.6 $\mu\text{m} \approx 2.07 \mu\text{m} \times \frac{12 \mu\text{m}}{9.48 \mu\text{m}}$ .
<sup>b</sup> Calculated by dividing MT growth velocity from Ref. [68] with average MT length, $k_{\text{ps}} \approx \frac{28 \mu\text{m}/\text{min}}{2.6 \mu\text{m}}$ .
<sup>c</sup> Based on electron microscopy data [8] which counted around 5400 non-kinetochore MTs in a cell, and EB1 measurements [68] which counted 62869 dots in 75 frames for MTs in 3 cells, giving an estimate of 280 growing MTs. Now, using the steady-state relationship between growing and pausing MTs from Eqs. (C8) and (C9) we calculate $k_{\text{cat}} = k_{\text{ps}} \frac{280}{5400 - 280}$ .
<sup>d</sup> Calculated using experimental data from [68]. Out of 280 growing MTs, 65% of the MT tracks in prometaphase are pointing toward the opposite pole, and 30% of MT tracks originate from zones up to 1.5 $\mu\text{m}$ away from the poles that can be ascribed to MTs nucleated at the pole. Consequently, the nucleation rate at the pole can be calculated from the estimate of the number of growing MTs nucleated at the pole as $\frac{k_p}{k_{\text{ps}}} = 280 \times 0.65 \times 0.30 = 54.6$ , from which we calculate $k_p$ .
<sup>e</sup> Calculated using experimental data from [68] where the number of MTs nucleated along pre-existing MTs growing toward opposite pole is $280 \times 0.65 \times 0.7 = 127.4$ . Using the parameters of our model we can estimate the number of growing MTs nucleated along one MT as $\frac{l_0}{k_{\text{ps}}} k_{nc}$ and multiplied by the number of pole MTs gives $N_p \frac{l_0}{k_{\text{ps}}} k_{nc} = 127.4$ , where
<sup>f</sup> Estimated as the ratio of the kinetochore area [48] to the cross-section area of the spindle [67].
<sup>g</sup> Estimated as force at the kinetochore, 300 pN [17], divided by growth velocity of kinetochore MTs, 1 $\mu\text{m}/\text{min}$ [35].
<sup>h</sup> Note that parameters $k_p$ and $p_{\text{att}}$ are relevant for the one-dimensional model but should be adjusted for more complex geometries.

**TABLE II.** Motor protein and crosslinker parameters.

| Parameter | Value | Description |
| --- | --- | --- |
| $c_m$ | 10 $\mu\text{m}^{-1}$ | Motor protein concentration<sup>a</sup> |
| $f_0$ | 5 pN | Motor protein stall force [72] |
| $v_0$ | 4 $\mu\text{m}/\text{min}$ | Motor protein velocity [35] |
| $c_c$ | 20 $\mu\text{m}^{-1}$ | Crosslinker concentration<sup>a</sup> |
| $\xi_c$ | 0.05 pN min / $\mu\text{m}$ | Crosslinker effective drag<sup>a</sup> |
| $N_{\text{CE}}$ | 20 | Nr. of motors at KC [73] |
| $f_{\text{CE0}}$ | 6 pN | KC motor stall force [74] |
| $v_{\text{CE0}}$ | 2.4 $\mu\text{m}/\text{min}$ | KC motor velocity<sup>a</sup> |

<sup>a</sup> Estimated.

### A. Motor proteins exert length-dependent forces that drive chromosome congression

In order to explore whether motor proteins can generate forces that drive chromosome congression, we reduce our model by setting the number of passive crosslinkers and kinetochore motor proteins to zero. In this regime we recalculate Eq. (18), which then simplifies to:

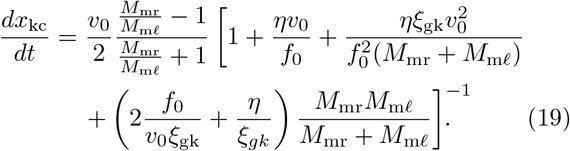

Here, it is evident that the kinetochore can move with velocity up to *v*_0_*/*2 and that the kinetochore moves toward the side with a larger number of motor proteins. We solve our model by numerically integrating Eq. (19), in which the number of motor proteins is calculated from Eqs. (7), (15) and (17).

For parameters relevant for human spindles(Tables I and II), the kinetochore, which was initially at a position close to the left pole, moves away from it and approaches the spindle equator zone in approximately 4 minutes (Fig. 2(a)). Based on this graph, we also estimate that the average velocity is about 1 μm*/*min. These predictions are consistent with the observed congression time, when compared with central chromosomes (Fig. 3f in Ref. [75]), as well as measured kinetochore velocity [26]. The quantitative agreement between theory and experiments suggests that the length-dependent forces generated by motor proteins can drive chromosome congression.

**FIG 2.**
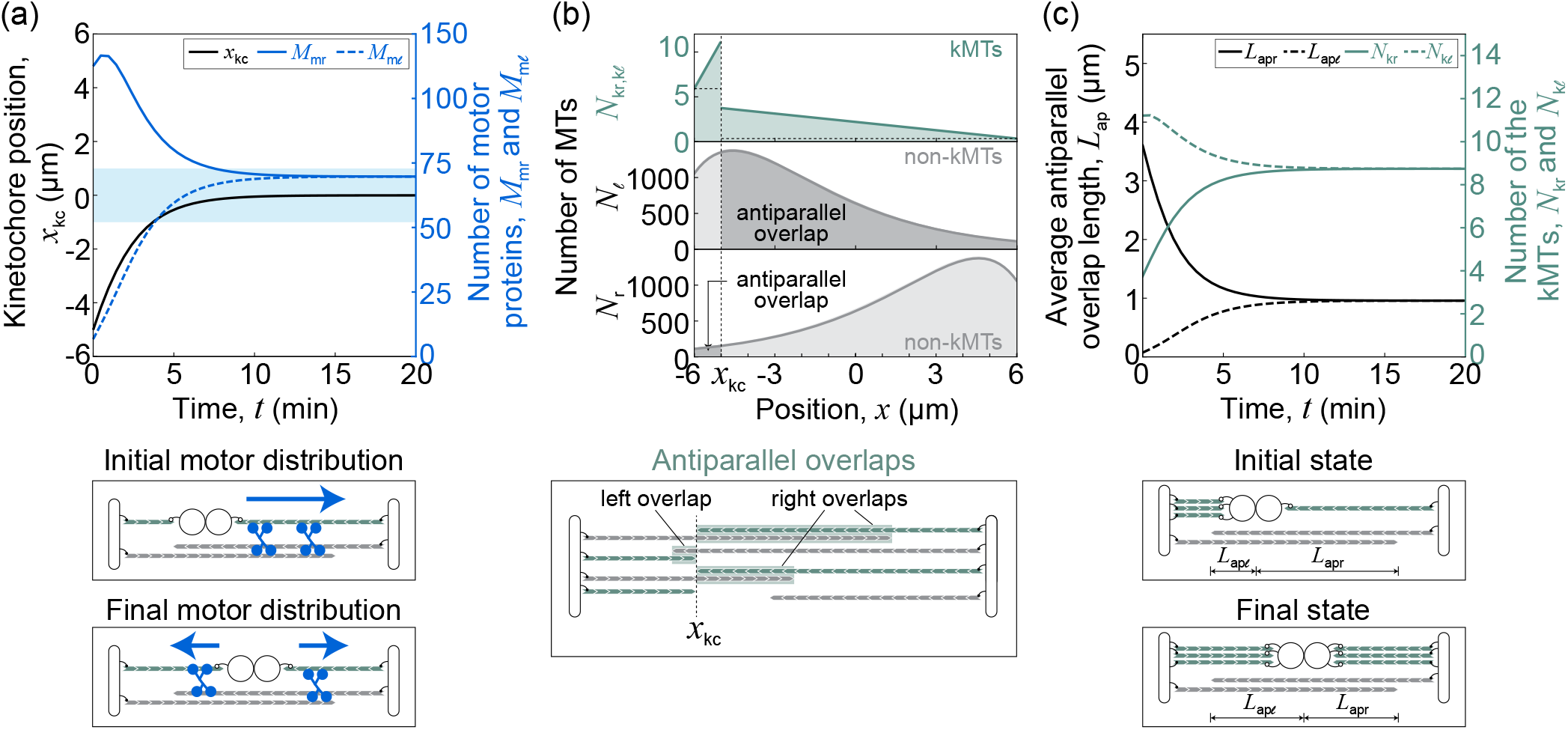
Motor proteins drive chromosome congression. (a) Solutions of the model for a case with only motor proteins, showing the time course of kinetochore position (black full line), and the number of motor proteins on the right (full blue line) and left (dashed blue line) for kinetochores initially at *x* = −5 μm. Blue area indicates the spindle equator zone. Schemes below panel (a) depict motor proteins (blue pictograms) and forces they exert (arrows) for initial and final kinetochore position. (b) Distributions of kinetochore MTs (top) and of non-kinetochore MTs (middle and bottom). The shaded regions indicate the fraction of MTs that form antiparallel overlaps with the kinetochore MTs. Number of right kinetochore MTs, at positions right from *x*_kc_ = −5 μm is calculated as *N*_kp_ + *N*_kn_(*x*), whereas number of left kinetochore MTs, at positions left from *x*_kc_ is calculated analogously. Analytical expressions for MT distributions on the left side *N*_kp*ℓ*_, *N*_kn*ℓ*_(*x*) and *N*_*ℓ*_ are given in Eqs. (D5), (D7) and (C10), respectively. Scheme below panel (b) depicts antiparallel overlaps for kinetochores at position *x*_kc_. (c) Time course of average antiparallel overlap length (black lines) and number of kinetochore MTs (green lines) for kinetochore positions as in panel (a). Solid lines represent values for the right side of the model, while dashed lines represent values for the left side. Schemes below panel (c) depict number of attached kinetochore MTs and the average antiparallel overlap length for initial and final kinetochore position. The parameters are given in Tables I and II, except for *c*_c_ = 0 and *N*_CE_ = 0.

**FIG 3.**
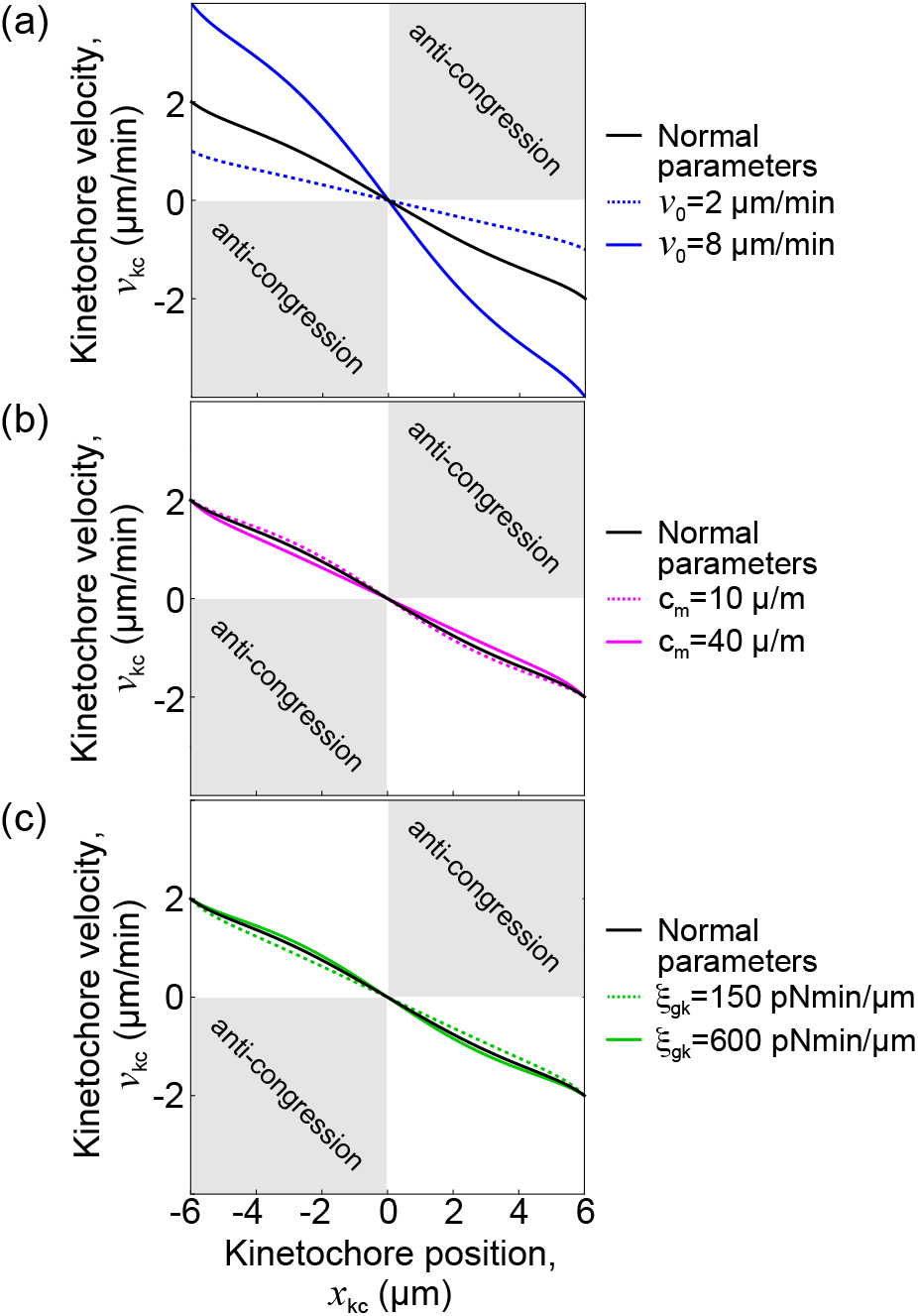
Kinetochore velocities as a function of the kinetochore position for original parameters, as well as two times greater and lower motor protein velocity without a load (a), density of motor proteins (b) and MT-kinetochore effective drag coefficient (c). The changed parameters are given in the legend. The remaining parameters are given in Tables I and II, except for *c*_c_ = 0 and *N*_CE_ = 0.

Kinetochore movement is driven by motor proteins that accumulate in greater numbers on the farther pole side than on the closer one (Fig. 2(a), blue lines). Because the number of motor proteins depends on antiparallel MT overlaps, we plot the distributions of kinetochore and non-kinetochore MTs for kinetochores at the initial position *x*_kc_ = −5 μm (Fig. 2(b)). Our calculations show that the number of kinetochore MTs increases linearly with proximity to the kinetochore. By comparing the left and right kinetochore MT distributions, we find that there are more kinetochore MTs on the closer pole side than on the farther one. The number of non-kinetochore MTs has a maximum in the vicinity of the respective pole and decreases in farther positions. By visualizing the fraction of non-kinetochore MTs that form antiparallel overlaps with kinetochore MTs, it can be seen that there is a substantially larger overlap region on the farther pole side as compared to the closer pole side (Fig. 2(b) middle and bottom). This difference is the main reason for a larger average antiparallel MT overlap length and consequently a greater number of motor proteins on the farther pole side.

In order to understand how the number of motor proteins changes during chromosome congression, we explore the time course of the average length of antiparallel overlap and the number of kinetochore MTs, as these two quantities directly influence the number of motor proteins (see Eq. (7)). We find that in the initial kinetochore position, the number of kinetochore MTs on the proximal pole side is almost three times greater, but because of the large difference in the antiparallel overlap lengths on the proximal and farther pole sides, the number of motor proteins is greater on the farther pole side (Fig. 2(c)). As kinetochores approach the final position they slow down and the number of kinetochore MTs and antiparallel overlaps become equal on both sides.

In our model, motor proteins generate forces that are directed toward the equatorial plane and thus drive chromosome congression. To visualize the centering efficiency, we plot the velocity of kinetochores as a function their position (Fig. 3). We show effects of both increasing and decreasing, by factor of 2, three relevant parameters: motor protein velocity without a load, density of motor proteins, and MT-kinetochore effective drag coefficient. We observe that the kinetochore velocity is oriented toward the center regardless of its position or changes in parameters. For kinetochores near the poles, the velocity magnitude is close to *v*_0_*/*2, decreasing as they approach the center, where it eventually reaches zero.

Furthermore, we explore kinetochore movement for different values of average MT length. As expected, there is no congression for very short MTs, because there are no kinetochore MTs on the farther pole side (Fig. 4 (a) case 1), but congression proceeds normally when the MTs are long enough to reach the kinetochore from the farther pole side (Fig. 4(a) cases 2 and 3).

**FIG 4.**
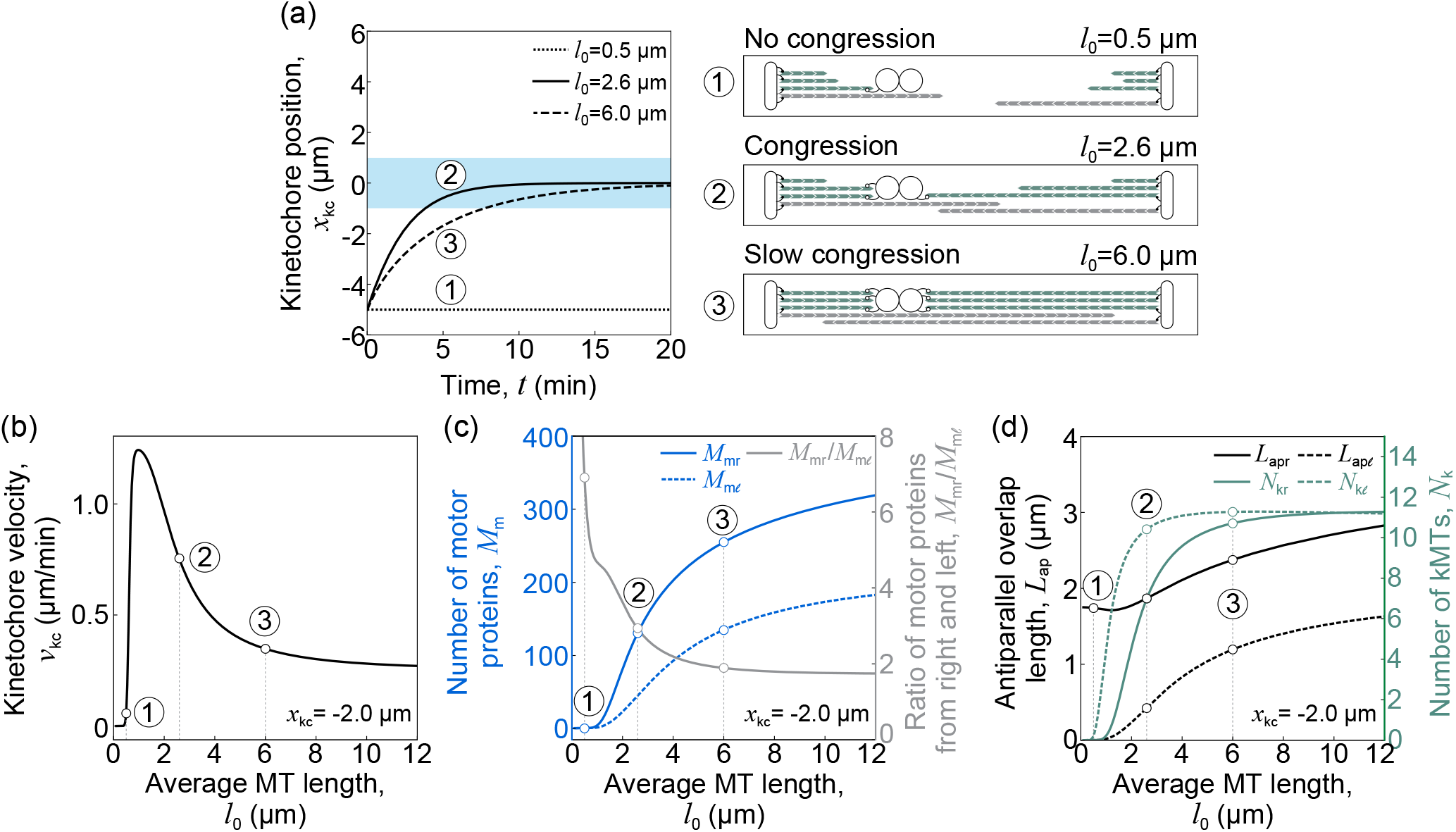
Congression velocity depends on average MT length. (a) (left) Solutions of the model showing time course of the kinetochore position for kinetochores initially at *x*_kc_ = −5 μm for different average MT lengths: cases with small MT length *l*_0_ = 0.5 μm (dotted line), normal MT length *l*_0_ = 2.6 μm (full line), large MT length *l*_0_ = 6 μm (dashed line) are denoted by encircled numbers 1-3, respectively. Light blue area indicates the spindle equator zone. (right) Schematic representations of the dependence of the numbers of kinetochore MTs corresponding to case 1, with no kinetochore MTs on the right kinetochore, case 2, with few kinetochore MTs on the right kinetochore, and case 3, with the same number of kinetochore MTs on both kinetochores. (b) Dependence of kinetochore velocities on average MT lengths. (c) Number of motor proteins at the right (full blue line) and left side (dashed blue line), and their ratio (gray line) as a function of the average MT length. (d) Average antiparallel overlap length of right (black line) and left (dashed black line) kinetochore MTs, and the number of kinetochore MTs (right y-axis) on the right (green line) and left (dashed green line) kinetochore as functions of the average MT length. In panels (b-d) kinetochores are at position *x*_kc_ = −2 μm. The encircled numbers 1, 2 and 3 show the values corresponding to the average MT lengths as in cases from panel (a). The remaining parameters are given in Tables I and II, except for *c*_c_ = 0 and *N*_CE_ = 0.

By calculating the velocity of the kinetochores as a function of the MT length, we find that for all lengths the velocity is directed toward the center (Fig. 4(b)). These calculations show that the kinetochore velocity reaches a maximum value for the average MT length around 1 μm. The kinetochore velocity has values comparable to biologically relevant values, which are around 0.5 μm*/*min [26], for an average MT length between 0.6 μm and 3.8 μm. To understand how the kinetochore velocity reaches the maximum value, we explore the dependence of the number of motor proteins on the average MT lengths (Fig. 4(c)). For small average MT lengths, the number of motor proteins is small. By increasing the MT length number of motor proteins and consequently their force increases, overcoming the chromosome drag friction, resulting in a greater kinetochore velocity (see Eq. (19)). On the other hand, a further increase in the average MT length results in a smaller ratio between the numbers of motor proteins on both sides, resulting in a decrease in the kinetochore velocity, even tough the number of motor proteins still increases.

In order to understand the relationship between the number of motor proteins and the average MT length, we explore changes in MT distributions. These distributions are represented by the length of antiparallel overlaps and the number of MTs attached to each kinetochore, the product of which is proportional to the number of motor proteins (see Eq. (7)). For small average MT lengths, we find a significant difference in length between left and right overlaps (see Fig. 4(d) for *l*_0_ below 2 μm), which remains for different average MT length, even though both lengths increase. This difference is the main reason for a greater number of motor proteins on the farther pole side. The number of kinetochore MTs is smaller on the farther pole side for smaller average MT lengths, but above 6 μm their difference becomes negligible. The relative difference in the number of kinetochore MTs on both sides is smaller as compared to the relative difference of the antiparallel overlap lengths, and thus the number of motor proteins is greater on the farther pole side.

### B. Forces exerted by passive crosslinkers impair chromosome congression

In the previous section, we showed that motor proteins distributed along antiparallel regions drive chromosome congression. Here, we explore to what extent passive crosslinkers accumulating in parallel regions affect chromosome congression. Unlike motor proteins, passive crosslinkers cannot generate active forces by themselves, and, for this reason, one can assume that passive crosslinkers cannot have an active role in chromosome congression. However, this reasoning changes when passive crosslinkers act in combination with active processes that generate directed movement. In our case, the directed movement of bridging MTs, poleward flux, drives the movement of kinetochore MTs as they are connected by passive crosslinkers. Thus, we explore the positioning of the kinetochores in our system in the presence of passive crosslinkers, but without motor proteins on the kinetochore MTs.

To study the role of crosslinkers in chromosome congression, we analyze the kinetochore velocity in the limit where the number of motor proteins and kinetochore motor proteins goes to zero (*M*_mr,m*ℓ*_ = 0 and *N*_CE_ = 0), for which Eq. (18) simplifies to:

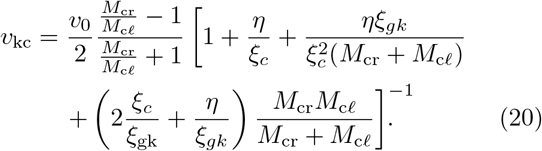

This equation shows that kinetochores move toward side with greater number of passive crosslinkers, which is similar to kinetochore movement driven by motor proteins (see Eq. (19)). Because passive crosslinkers accumulate in parallel overlaps and motor proteins in antiparallel overlaps, we expect their distributions to differ and thus differently affect chromosome congression.

Our numerical calculations show that for biologically relevant MT lengths the kinetochores do not approach the spindle center. Instead, they asymptotically approach one of two stable points located near poles, where the point to which the kinetochores approach depends on their initial position. (Fig. 5(a)).

**FIG 5.**
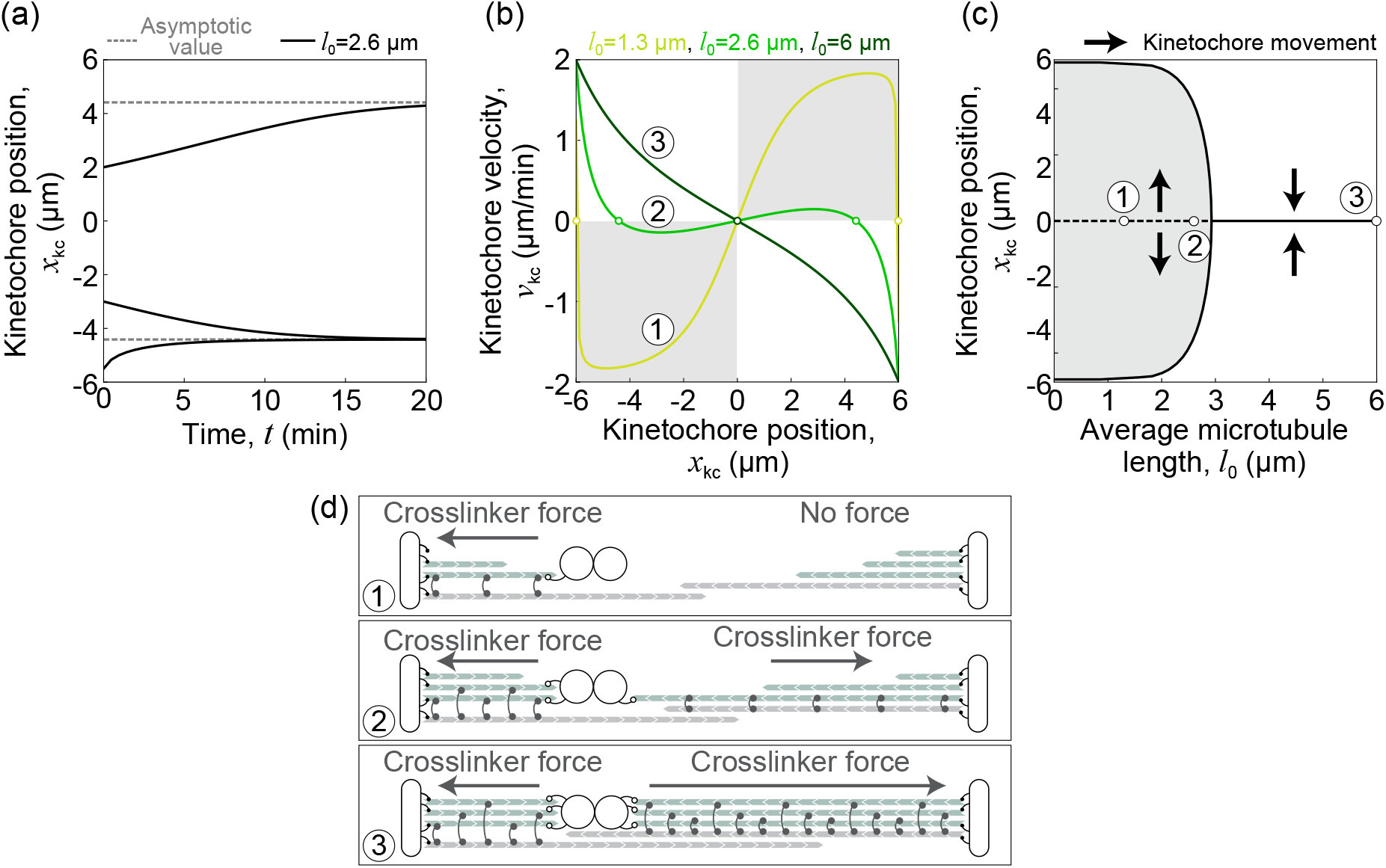
Passive crosslinkers produce forces that can oppose chromosome congression. (a) Time courses of the kinetochores position (black lines) driven by crosslinkers for the kinetochore starting at three different positions, *x*_*kc*_ = −5 μm, *x*_*kc*_ = −3 μm and *x*_*kc*_ = 2 μm. The kinetochores approach one of two asymptotic positions (gray dashed lines). (b) Dependence of the kinetochore velocity on the kinetochore position along the MT spindle for three different average MT lengths: short MTs *l*_0_ = 1.3 μm (case 1, pear color line), normal MT lengths *l*_0_ = 2.6 μm (case 2, green line) and long MTs *l*_0_ = 6 μm (case 3, dark green line). The gray areas represent the choice of positions and velocities for which the kinetochores will move toward the closer pole. White dots represent stable fixed points for different average MT lengths. (c) Positions of the stable (black full line) and unstable (black dashed line) fixed points for different average MT lengths. The gray area represents the choice of positions and MT lengths for which the kinetochores will move toward the closer pole. The kinetochore movement direction is indicated by the black arrows. (d) Schematic depiction of forces exerted by crosslinkers (black) on kinetochores (white) for short MTs (case 1), intermediate MTs (case 2) and long MTs (case 3). Force magnitude and direction on each kinetochore are indicated by an arrow. In panels (a)-(c) the unchanged parameters are given in Tables I and II, except for *c*_m_ = 0 and *N*_CE_ = 0.

To further explore the kinetochore movement, we show the dependence of kinetochore velocity on its position for different values of the average MT length (Fig. 5(b)). For average MT lengths *l*_0_ = 1.3 μm and *l*_0_ = 2.6 μm there are two stable points placed symmetrically with respect to the unstable point located in the center. We refer to these three points as fixed points. For larger average MT lengths, *l*_0_ = 6μm, the central location changes stability and becomes a unique stable fixed point that the kinetochores approach irrespective of its initial position.

To describe the system in the vicinity of the central fixed point, we explore the stability of this point (Fig. 5(c)). By numerically calculating Eq. (E1), we find that the system undergoes a supercritical pitchfork bifurcation with a critical average MT length of *l*^*\**^ ≈ 3 μm. In this case, above the critical MT length there is only one stable fixed point in the center. Below the critical MT length central fixed point becomes unstable, and the transition in stability is accompanied by the appearance of two stable fixed points.

To gain an intuitive explanation why the fixed point in the center changes its stability we depict distributions of MTs for three different average MT lengths (Fig. 5(d)). For small average MT lengths MTs do not reach kinetochores from the far pole side, whereas from the near pole side MTs reach the kinetochore (case 1). In this case passive crosslinkers accumulate within the parallel overlap at the near pole side only and thus generate off-centering force. As average MT length increases, MTs reach the kinetochore from the far pole side and crosslinkers accumulate at both sides, resulting in similar forces on both kinetochores (case 2). For large average MT lengths, the number of kinetochore MTs becomes similar on both sides, thus a longer overlap on the farther pole side accumulates more crosslinkers producing a centering force (case 3).

### C. The collective forces of motor proteins and crosslinkers determine the direction of chromosome movement

In previous subsections, we separately explored the influence of motor proteins and crosslinkers on chromosome congression, whereas in real biological systems they work together, and thus we explore their combined contribution. For biologically relevant parameters, the kinetochores that were initially displaced from the spindle center approach it in several minutes (Fig. 6(a)). This outcome occurs regardless of initial conditions, for kinetochores close to the pole and those closer to the center. The dynamics of this process are similar to those in the case without crosslinkers (Fig. 2(a)), implying that motor proteins generate dominant forces during chromosome congression.

**FIG 6.**
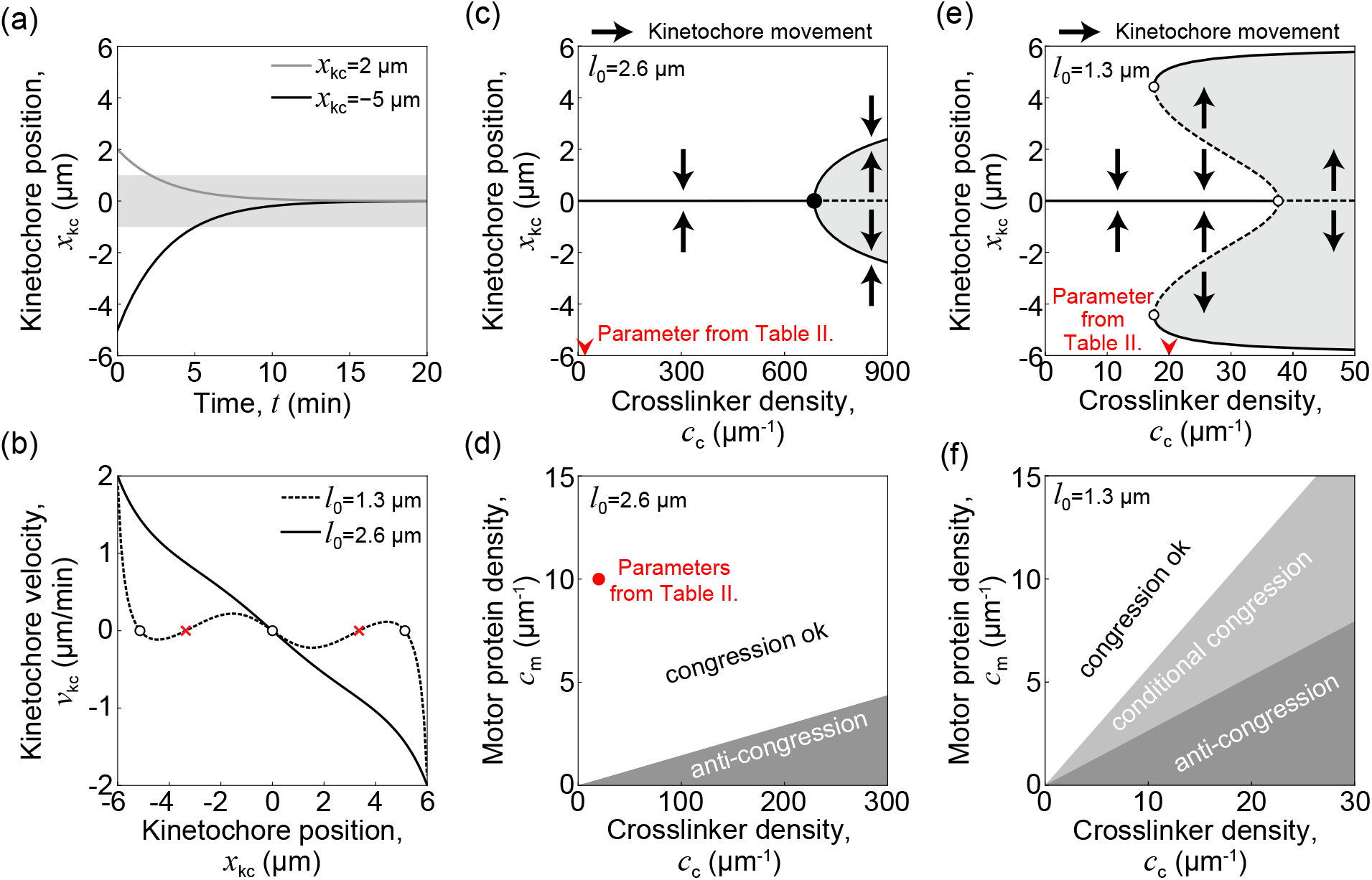
Interplay of motor protein and crosslinker forces regulates the effectiveness of chromosome congression. (a) Solutions of the model showing time course of kinetochore position for kinetochores initially at *x* = 2 μm (gray line) and *x* = −5 μm (black line). Blue area indicates the spindle equator zone. (b) Dependence of the kinetochore velocity on the kinetochore position for two different MT lengths *l*_0_ = 3 μm (black line) and *l*_0_ = 1.5 μm (gray line). The dots represent the stable fixed points, whereas the red “x” symbol denotes the unstable fixed points. (c) The position of the stable (black full line) and unstable (black dashed line) points for different average crosslinker densities and MT length *l*_0_ = 2.6 μm. The arrow on the x-axis represents the crosslinker density *c*_*c*_ given in Table II. The kinetochore movement direction is indicated by the black arrows. (d) Phase diagram showing a region where congression occurs for different crosslinker and motor protein densities for MT length *l*_0_ = 2.6 μm. The transition line occurs at *c*_m_ ≈ 0.019 *c*_c_. The red dot represents the density values given in Table II. (e) The position of the stable (black full line) and unstable points (black dashed line) for different average crosslinker densities and MT length *l*_0_ = 1.3 μm. (f) Phase diagram showing a region where congression occurs for different crosslinker and motor protein densities and MT length *l*_0_ = 1.3 μm. The two transition lines occur at *c*_m_ ≈ 0.67 *c*_c_ and *c*_m_ ≈ 0.29 *c*_c_. The kinetochore movement direction is indicated by the black arrows. The remaining parameters are given in Tables I and II, except for *N*_CE_ = 0.

Our results show that average MT length plays an important role in chromosome movement. To explore how the velocity of kinetochores depends on their position when both motor proteins and crosslinkers are present, we plot this dependence for two different values of average MT lengths (Fig. 6(b)). For parameters as in Tables I and II, all velocities are directed toward the stable central position. However, for smaller average MT lengths, there are three stable fixed points and two unstable fixed points located between neighboring stable fixed points, leading to increased complexity in the kinetochore movement.

Next, we explore the stability of the central position for different values of crosslinker density (Fig. 6(c)). For a wide range of crosslinker densities, the central position is the only fixed point and all kinetochores move toward it, irrespective of the starting position. When the concentration exceeds a critical value, the central location becomes unstable, and two new stable fixed points appear symmetrically with respect to the center, which is a signature of supercritical pitchfork bifurcation. Further, we explore the parameter space by studying kinetochore movement in the vicinity of the central fixed point, as in Appendix E. We find that the critical crosslinker density, for which the central fixed point changes its stability, is linearly proportional to the density of motor proteins (Fig. 6(d)). For values below the critical crosslinker density, congression proceeds normally, while for values above it chromosomes move away from the center.

Motivated by the complex kinetochore dynamics for smaller average MT lengths (Fig. 6(b)), we explore the transitions that occur in this region of parameter space. Using crosslinker density as a control parameter, we plot the position of stable and unstable fixed points (Fig. 6(e)). The results show a subcritical pitchfork bifurcation, where an unstable fixed point becomes stable, and two new unstable fixed points emerge as the crosslinker density decreases. For the same crosslinker densities there are two stable fixed points and they coexist with the unstable fixed points, on both sides of the bifurcation.

Depending on the number of stable fixed points, we identify three distinct regions in the parameter space (Fig. 6(f)). These results suggest that, when there is more than one stable fixed point, congression fails and kinetochores can be stably located both near the poles and at the center.

### D. Plus-end-directed kinetochore motor proteins assist chromosome congression

In addition to motor proteins and crosslinkers, our model also describes kinetochore motor proteins that contribute to chromosome congression by transporting chromosomes along MTs toward their plus-ends. In order to isolate the influence of kinetochore motor proteins on chromosome congression, we set the number of motor proteins and crosslinkers in Eq. (18) to zero, yielding the kinetochore velocity:

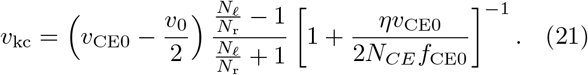

Here, we see that the kinetochore velocity direction depends on the difference between the kinetochore motor protein velocity without a load and the non-kinetochore MT flux velocity. This suggests that kinetochore motor proteins driven chromosome congression is possible when these motors are faster than the poleward flux, providing an explanation for different directions of chromosome movement in Fig. 7(a). Based on Eq. (21), we also find that the kinetochore velocity depends on the ratio of the number of left and right non-kinetochore MTs, whereas calculations in Appendix F show under which conditions kinetochore motor proteins generate anti-congression.

**FIG 7.**
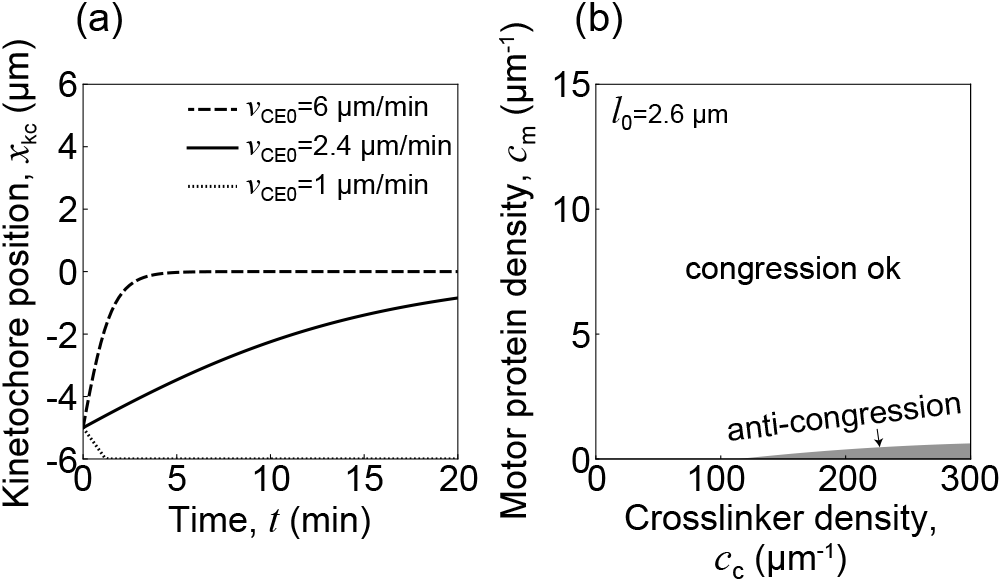
Chromosome congression with kinetochore motor proteins alone and in combination with other proteins. (a) Solutions of the model showing the time course of the kinetochore position driven by kinetochore motor proteins affects the kinetochore MTs starting at *x*_*kc*_ = −5 μm for different velocities without a load of the kinetochore motor proteins. (b) Phase diagram showing a region where congression occurs for different crosslinker and motor protein densities. The remaining parameters are given in Tables I and II.

To explore how kinetochore motor proteins work together with motor proteins and crosslinkers to promote congression, we plot a phase diagram by varying the crosslinker and motor protein densities (Fig. 7(b)). These calculations reveal that chromosome congress in a large fraction of parameter space. In comparison with Fig. 6(d), we observe that the region of parameter space for which chromosomes congress increases in the presence of kinetochore motor proteins, suggesting that they promote chromosome congression. Taken together, our theory shows that length-dependent poleward flux drives chromosome congression for a broad range of biologically relevant parameters.

## IV. DISCUSSION

Our model highlights the importance of forces proportional to MT overlap lengths in chromosome congression. Our calculations show that motor proteins, which accumulate in antiparallel overlaps, at long kinetochore MTs produce forces that overcome those generated on the shorter kinetochore MTs, resulting in chromosome congression. Conversely, passive crosslinkers produce forces that pull the chromosomes toward the closer pole, opposing congression. Crucially, the forces generated by the motor proteins are large enough to overcome the forces that oppose congression. Our calculations also show that plus-end-directed motor proteins at kinetochores assist congression. Thus, our model provides a suitable tool for studying forces relevant for chromosome congression and reproduces experimentally measured kinetochore velocities, including duration of chromosome congression.

It has been shown that regulation of length of MTs attached to kinetochores is essential for optimal congression velocity, where MT dynamics is controlled by motor protein Kif18A, which accumulates in a length dependent manner [31, 76, 77]. Our model provides an alternative explanation, where kinetochore MTs form overlaps with non-kinetochore MTs and thus exert forces that are length dependent (Fig 2). These two mechanisms do not oppose each other, and thus they can work together, but future experiments will clarify the contribution of each individual mechanism.

In experiments in cells with altered concentrations of CENP-E proteins, chromosomes can be found in the metaphase plane, as well as near the poles [26]. Such states can persist for long periods of time, even after inhibiting Aurora B (which detaches kinetochore MTs at low inter-kinetochore tension), suggesting that there are several stable locations along the spindle where kinetochores tend to accumulate. Our model predicts that a regime with multiple stable points can appear. For example, in spindles with small average MT length stable points in the center and near the pole coexist, and kinetochores approach one of them depending on the initial position (Fig. 6(b), (e) and (f)). A close comparison of our theoretical predictions and experiments that quantify the distribution of MTs attached to kinetochores will provide a deeper insight into the mechanisms that help in the correction of these erroneous states.

Our model describes movement of one chromosome, implying that chromosomes move independent of each other. Study of chromosome movement during metaphase shows that chromosomes typically move independent of each other, but in case of neighboring chromosomes there is a certain correlation in the movement of chromosomes [78]. Based on this observation it was proposed that the correlation is a consequence of interaction between kinetochore MTs of neighboring chromosomes. In our mechanism of chromosome congression, forces at kinetochore MTs arise from interactions with neighboring MTs. Thus, extending the model to include multiple chromosomes and interaction between their kinetochore MTs could result in correlated movement of neighboring chromosomes. Future studies, theoretical and experimental, will reveal to which extent chromosome movement is correlated during congression.

During mitosis MTs from one side can attach to either kinetochore and occasionally form erroneous, merotelic or syntelic attachments [79, 80]. These types of errors are typical in tumor cells, so it would be interesting to explore the formation and correction of such attachments by a theoretical model. Our model could be generalized by describing MT attachment to kinetochores from both sides, as well as the observed tension-dependent MT detachment from kinetochores [63, 81] and Aurora B activity [82–84]. Such a model could provide a deeper understanding of the correction of different types of erroneous attachments.

Our model describes chromosome congression, but the framework we propose can also be extended to metaphase. For instance, it may offer insights into the centering forces that govern kinetochore oscillations. In existing models, the centering force is typically attributed to polar ejection force [54, 56, 57]. Future studies will help determine which centering force plays the dominant role in driving kinetochore oscillations.

In conclusion, we introduced a model that describes the most important forces that appear during chromosome congression, and therefore it represents a powerful tool for studying this biological process. This model relies on length-dependent poleward flux and describes MT distributions in mean-field approximation, which allows us to systematically explore the parameter space and distinguish the contributions of different mechanisms.

## V. ACKNOWLEDGMENT

We thank Kruno Vukušić, Iva Tolić and Patrick Meraldi for valuable discussions and help in comparing experiments and theory, also Maja Novak and Shane Amadeus Fiorenza for comments on the manuscript and other members of the N.P and I.M.T. research groups for their feedback. This work was funded by the European Research Council (ERC Synergy Grant, GA number 855158, granted to N.P.) and co-funded by the Croatian Science Foundation (project IP-2019-04-5967 granted to N.P.)

## Appendix A: Equations of movement of left and right kinetochore

In the main text of the paper, we focus on equations describing the movement of the center of mass of the sister kinetochores. Here, we present the full equations describing the movements of the left and right kinetochore, and how to get the center of mass equations. The equations of movement of the left and right kinetochore are given as:

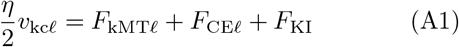

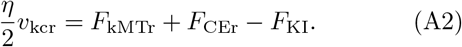

Here, 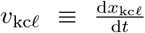 and 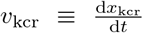 denote the velocities of the left and right kinetochore, respectively. *F*_KI_ denotes the force of interaction between two kinetochores. Inserting these two equations into the definition of the position of the center of mass of sister kinetochores 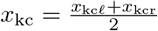, we get Eq. (1), rewritten as:

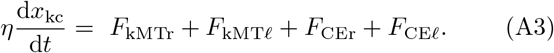

In the paper, we approximate each of these forces as a function of the position of the center of mass of the left and right kinetochores only and not the positions of the individual kinetochores. In order to be able to calculate the velocity of the kinetochores it is necessary to provide further definitions for the left and right side functions. Kinetochore MT growth velocities, defined as

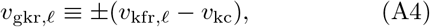

are calculated as

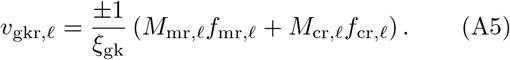

Here and in these appendices the upper sign (“+” or “-”) corresponds to the right side index “r”, where the lower sign corresponds to the left side index “*ℓ*”. The average force per motor protein *f*_mr,*ℓ*_ for both sides is calculated as

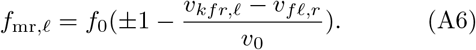

Similarly, the average drag force per crosslinker is calculated as

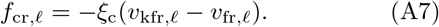

Additionally, the forces due to kinetochore motor proteins stepping toward right or left are calculated as

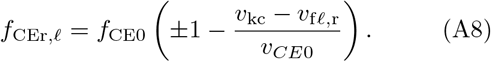

Where we redefine the average kinetochore motor protein force, as *f*_CE,pp*ℓ*_ = *f*_CE,apr_ = *f*_CEr_ and *f*_CE,ppr_ = *f*_CE,ap*ℓ*_ = *f*_CE*ℓ*_. The equations governing non-kinetochore MT distributions for the left and right sides are identical to those in Eqs. (9) -(12) by form. However, in Eq.(11), the number of pole MTs at position *x* pointing toward the left, *N*_pr_(*x*), is connected to the number of pole MTs at position *x* pointing toward the right as *N*_p*ℓ*_(*x*) = *N*_pr_(−*x*). The equations for the distributions of kinetochore MTs on the left and right kinetochore differ in one term due to the kinetochore velocity affecting the attachment:

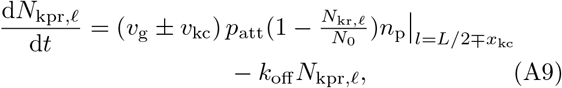

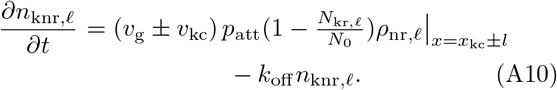

The number of kinetochore MTs on the right and left kinetochores is calculated as:

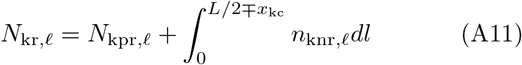

and the average parallel and antiparallel overlap lengths as:

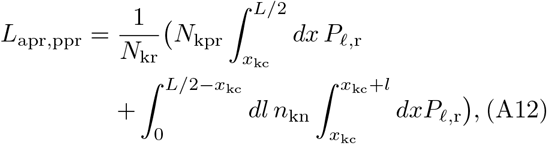

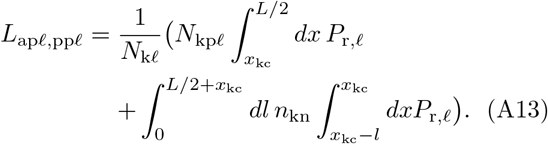

Here we presented the equations for the left and right sides corresponding to Eq. (1) - Eq. (17)

## Appendix B: Calculating the kinetochore velocity

In order to calculate the kinetochore velocity, *v*_kc_, we start with the equations for the balance of forces on the kinetochore,

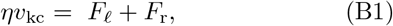

and by applying Eqs. (6) and (8) we get:

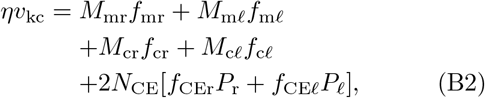

where we applied the redefinition of the kinetochore motor proteins forces as in Eq. (A8). We proceed by inserting the force-velocity expression from Eqs. (A6) - (A8). The sum of the kinetochore-associated motor protein forces is:

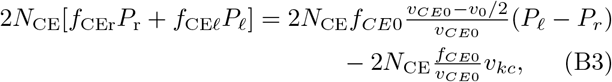

where we used *v*_*fr*,*ℓ*_ = *±v*_0_*/*2 and *P*_*ℓ*_ + *P*_*r*_ = 1. The sum of the forces due to motor proteins and crosslinkers on the kinetochore MTs follows as:

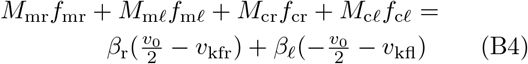

where we used a shorthand notation 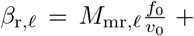. To find the kinetochore MT flux velocities, *v*_kfr,*ℓ*_, we use the equations for the growth of kinetochore MTs, Eq. (A5), and by inserting the force-velocity expression from Eq. (A6) - (A8) we get:

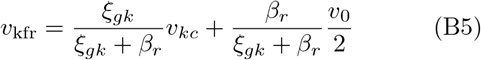

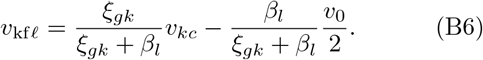

Inserting these equations into (B4) yields the equation that shows the kinetochore velocity as a function of the total number of motor proteins, crosslinkers and kinetochore-associated motor proteins:

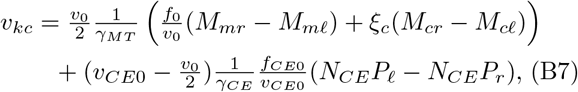

where we introduced additional shorthand notations: 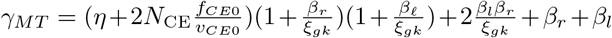 and 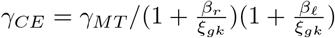.

## Appendix C: Calculating the non-kinetochore microtubule distributions

Here we calculate the non-kinetochore MT distributions in a steady state limit. The stationary equations for distributions of non-kinetochore MTs that point rightwards, calculated from Eq. (9) - Eq.(12), are given as:

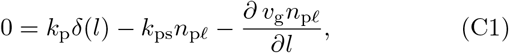

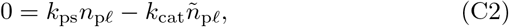

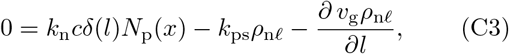

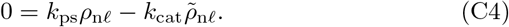

We solve the equations Eq. (C1) and Eq. (C3) in two steps. First, we find the solutions of the homogeneous part, which for these two equations have the same form with different integration constants, 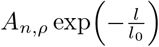, where

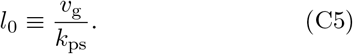

By taking into account the inhomogeneous terms we calculate the integration constants yielding the stationary distributions of non-kinetochore MTs extending from the pole and those nucleated along pre-existing MTs:

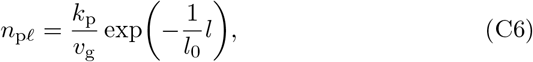

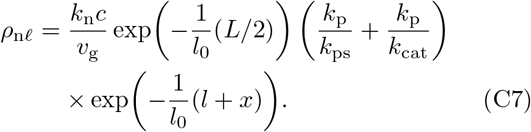

The solutions of Eq. (C2) and Eq. (C4) provide a simple relationship between the number of growing and pausing MTs:

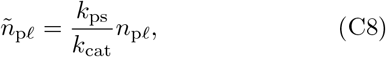

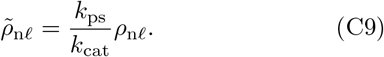

Based on the calculated analytical expressions for MT distributions we can obtain the number of MTs at any position along the spindle axis. The MTs that cross the position *x* are those with the minus-end and plus-end on opposite sides of that position. Thus, the calculation for the number of MTs pointing rightwards is given as:

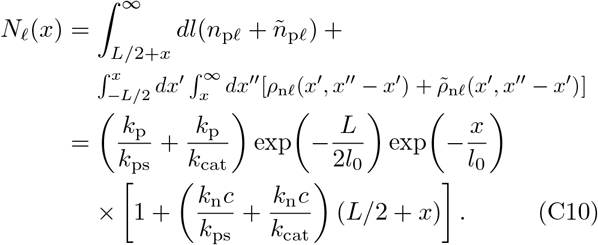

The distributions for MTs pointing leftward are given by substituting *x* → −*x* in Eq. (C7), Eq. (C9) and Eq. (C10), whereas Eq. (C6) and Eq. (C8) depend on MT length only, and thus are identical for both directions.

## Appendix D Calculating the kinetochore microtubule distributions

We calculate the kinetochore MT distributions in a steady state limit. We also neglect the kinetochore velocity, *v*_kc_, since it is much smaller than the MT growth velocity *v*_g_. With these approximations Eqs. (13) and (14) simplify to:

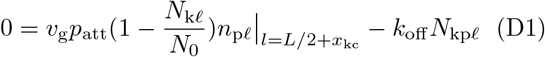

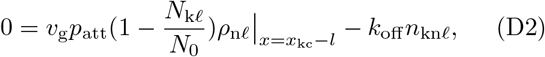

Note that these equations describe MTs attached to the left kinetochore. These coupled algebraic equations, Eq. (D1) and Eq. (D2), link the distributions of kinetochore MTs that extend from the pole and along pre-existing MTs, yielding a linear relationship with a coefficient that depends on the position along the spindle:

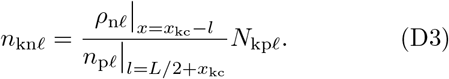

Inserting this result in the equation for the total number of MTs at the left kinetochore, Eq. (15), and using the fact that 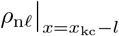 does not depend on *l*, it follows that

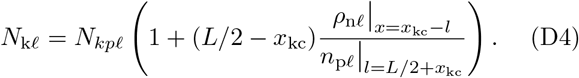

Combining this result with Eqs. (D1) and (D2), the number of left kinetochore MTs that reach the pole is given as

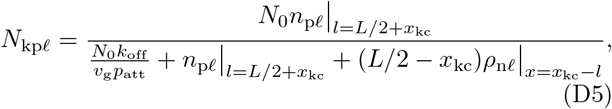

whereas the length distribution of MTs that do not reach the pole is given as

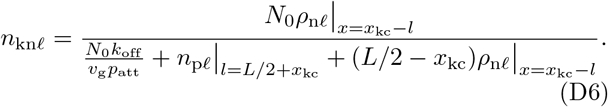

The number of kinetochore MTs nucleated along pre-existing MTs that reach position *x* is calculated as:

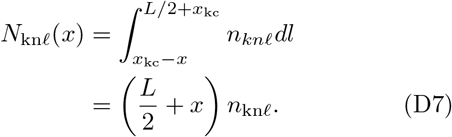

In particular, the total number of kinetochore MTs nucleated along pre-existing MTs is given by *N*_kn*ℓ*_(*x*_kc_).

The kinetochore MT distributions on the right kinetochore follow by substituting *x* → −*x* and *x*_kc_ → −*x*_kc_ in Eqs. (D5) - (D7), as well as by using the right side MT distributions instead of the left.

## Appendix E: Stability analysis for kinetochores in the vicinity of the spindle equator

In order to explore the stability of the positioning of kinetochores around the spindle center, we calculate the kinetochore velocity direction in the vicinity of the spindle center. From Eq. (C10) follows a mirror symmetry with respect to the spindle center for the probability of motor protein attachment for the left and right side *P*_*l*_(*x*) = *P*_*r*_(−*x*). Due to this symmetry, there is also a mirror symmetry for the motor protein and crosslinker numbers for the left and right side *M*_mr_(*x*) = *M*_m*ℓ*_(−*x*) and *M*_cr_(*x*) = *M*_c*ℓ*_(−*x*). By using these symmetries to calculate the kinetochore velocity, from Eq. (13) it follows that the velocity is antisymmetric with respect to the spindle center *v*_kc_(*x*) = −*v*_kc_(−*x*), and it is equal to zero at the spindle center. We calculate the kinetochore velocity in the vicinity of the spindle center, at position *x*_kc_ = *ϵ*, which is small compared to the typical MT length |*ϵ*| ≪ *l*_0_. Taylor expansion of *v*_kc_ as a function of the position *x* at the point *x* = 0 up to the first term gives 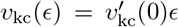. Here and throughout the text, the prime symbol ^*′*^ denotes the first derivative of a function with respect to *x*. By taking the first derivative of Eq. (13) at *x* = 0 and by using the mirror symmetries of the motor proteins, crosslinkers and the motor protein attachment probabilities we find:

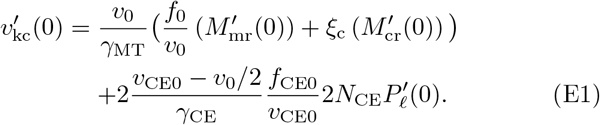

This equation provides straightforward information about the stability of the kinetochore positioning, where for negative values of the velocity derivative the kinetochore is at a stable position.

## Appendix F: Movements of kinetochore due to kinetochore motor proteins

In section III D it was shown that the kinetochore velocity is oriented toward the center under the influence of the kinetochore motor proteins. However, it is possible to find a choice of parameters for which the kinetochore velocity is not oriented toward the center. We proceed from Eq. (E1) and calculate:

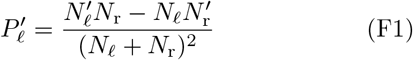

Next, we substitute the expressions for *N*_*ℓ*_ and *N*_r_ calculated from equation (C10) at position *x*_kc_ = 0 :

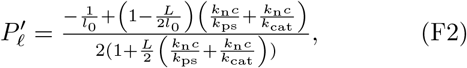

From this equation, we see that when no nucleation along pre-existing MTs is present or for *l*_0_ *< L/*2 we get 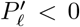. In this case, if *v*_CE_ *> v*_0_*/*2, the contributions of kinetochore motor proteins to the force driving the movements of chromosomes are centerward. On the other hand, if nucleation along pre-existing MTs satisfies 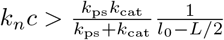 and if *l*_0_ *> L/*2 we have 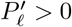. In this case, if *v*_CE_ *> v*_0_*/*2, the contributions of kinetochore motor proteins to the force driving the movements of the chromosomes are directed toward the poles in the region around the spindle center.

## Notes

### Competing Interest Statement

The authors have declared no competing interest.

### Summary of Updates

This version of the manuscript has been revised to make it more approachable for non-experts by adding schemes in Fig. 1. The clarity of certain statements were improved, which led to revising the title and two paragraphs. Additional explanation was provided for the one-dimensional mean-field approach by elaborating on the biological context of the model.

